# *Bacillus clarus* sp. nov. is a new *Bacillus cereus* group species isolated from soil

**DOI:** 10.1101/508077

**Authors:** Marysabel Méndez Acevedo, Laura M. Carroll, Manjari Mukherjee, Emma Mills, Lingzi Xiaoli, Edward G. Dudley, Jasna Kovac

**Author notes:** Corresponding Author: Jasna Kovac, 437 Rodney A. Erickson Food Science Building, University Park, PA, 16802. GenBank/EMBL/DDBJ accession number for the 16S rRNA of isolate PS00077AT^T^ is MH918154. WGS accession number for the draft genome of isolate PS00077AT^T^ is QVOD00000000. Two supplementary figures and one supplementary table are available with the online version of this paper.

## Abstract

*Bacillus cereus* group or *B. cereus* sensu lato (s.l.), is comprised of Gram-positive spore-forming, rod-like bacteria that are widespread in natural environments. Although the species in this group are known to be highly related in terms of phenotypic characteristics, they display different levels of pathogenicity. Biochemical assays are therefore considered to be insufficient for accurate taxonomic classification of *B. cereus* group species. To facilitate accurate taxonomic classification and associated prediction of pathogenic potential, we have conducted comparative genomic analyses of publicly available genome assemblies of *B. cereus* group isolates. Through that, we found that an isolate previously known as *B. mycoides* ATCC 21929 was sufficiently distant from valid and effective type strains to be considered a putative new species. We have conducted biochemical and bioinformatic characterization of strain ATCC 21929 that had been isolated from soil in Papua New Guinea. Strain ATCC 21929 most closely resembles *B. paramycoides* NH24A2^T^, producing ANIb and DDH values of 86.70% and 34.1%, respectively. Phenotypically, isolate ATCC 21929 does not possess cytochrome c oxidase activity, and is able to grow at a range of temperatures 15°C - 43°C and at a range of pH 6 - 9. With regards to fatty acid composition, this isolate has iso-C17:0 in highest abundance. We propose the strain ATCC 21929^T^ (=PS00077A^T^ = PS00077B^T^ = PSU-0922^T^ = BHP^T^) as a new species named *Bacillus clarus* sp. nov. to facilitate accurate taxonomic classification of *B. cereus* group isolates.

---

The *Bacillus cereus* group currently comprises 18 valid species, including *B. cereus* sensu stricto (s.s.) [1–3], *B. albus* [4], *B. anthracis* [5], *B. thuringiensis* [6], *B. mycoides* [7], *B. cytotoxicus* [8], *B. luti* [4], *B. mobilis* [4], *B. nitratireducens* [4], *B. pacificus* [4], *B. paramycoides* [4], *B. paranthracis* [4], *B*. [4], *B. pseudomycoides* [9], *B. toyonensis* [10], *B. tropicus* [4], *B. wiedmannii* [11], and *B. weihenstephanensis* [7]. In addition to these valid species, the group contains effectively published species (i.e., “B. gaemokensis” [12], “B. manliponensis *“* [13] and “B. bingmayongensis” [14]). Strains from the *B. cereus* group are commonly found as part of the plant and soil microbiome [15]. They can form spores, and they are rod-shaped and facultatively anaerobic [16]. Some *B. cereus* group isolates are important biocontrol agents (e.g., insecticidal *B. thuringiensis* isolates), while others can cause food spoilage or disease in animals or humans (e.g., anthrax, emetic and diarrheal gastrointestinal disease) [1, 15–17]. We have recently developed and published BTyper, a *B. cereus* group whole genome sequence-based comprehensive and rapid genotyping tool [18]. We used BTyper in conjunction with FastANI [19] to investigate genomic diversity of *B. cereus* group isolates for which assembled genomes had been made available through NCBI. We have identified strain BHP with RefSeq assembly accession GCF_000746925.1 as a putative new species based on average nucleotide identity BLAST (ANIb) and *in silico* DNA-DNA hybridization (DDH) values. The strain most closely resembled *B. paramycoides* NH24A2^T^, producing ANIb and DDH values of 86.70% and 34.1%, respectively. Strain BHP is also known under alias *B. mycoides* ATCC 21929 and has been reported previously as a producer of an antibiotic compound active against Gram-positive pathogens [20]. For this study, *B. mycoides* ATCC 21929 was obtained from ATCC for whole genome sequencing to verify its identity and for phenotypic characterization. Based on genomic and phenotypic dissimilarities of *B. mycoides* ATCC 21929 compared to valid and effective *B. cereus* group species, we propose the strain ATCC 21929^T^ as a type strain of a new species *B. clarus* sp. nov. *B. clarus* type strain is deposited in ATCC (=ATCC 21929^T^) and is also known under aliases BHP^T^, PS00077A^T^, PS00077B^T^ and PSU-0922^T^.

## Phylogenetic analyses

The genome of the proposed novel *B. cereus* group species *B. clarus* ATCC 21929^T^ (=BHP^T^) was downloaded from NCBI’s RefSeq database (RefSeq accession GCF_000746925.1, deposited by Los Alamos National Laboratory in 2014) (Pruitt 2007 Nucleic Acids Res). To confirm the identity of the isolate obtained from ATCC that was phenotypically characterized in this study, the strain was re-sequenced at the Penn State Department of Food Science as part of the FDA GenomeTrakr effort. The re-sequenced genome is accessible under NCBI accession number QVOD00000000. The RefSeq sequence GCF_000746925.1 was used in all further phylogenetic analyses reported in this paper.

To confirm that the isolate ATCC 21929^T^ was a member of the *B. cereus* group, its genome sequence was compared to a database of 16S rRNA sequences of other *B. cereus* group type strains (Table 1) using nucleotide BLAST (blastn) version 2.6.0 [21] and BTyper version 2.2.2 [18]. BTyper version 2.2.2 [18] was used to extract 16S rDNA gene sequences from the genome of the proposed novel species *B. clarus* ATCC 21929^T^ and the 18 published *B. cereus* group species. MUSCLE version 3.8.31 [22, 23] was used to construct an alignment of all 19 16S rDNA genes. RAxML version 8.2.11 [24] was used to construct a maximum likelihood (ML) phylogeny based on the 16S rDNA gene alignment, with the General Time-Reversible (GTR) nucleotide substitution model under the Gamma model of rate heterogeneity and 1,000 bootstrap replicates. FigTree version 1.4.3 (http://tree.bio.ed.ac.uk/software/figtree/) and Graphic for Mac 3.1 were used to annotate the resulting phylogenetic tree (Fig. 1).

**Figure 1.**
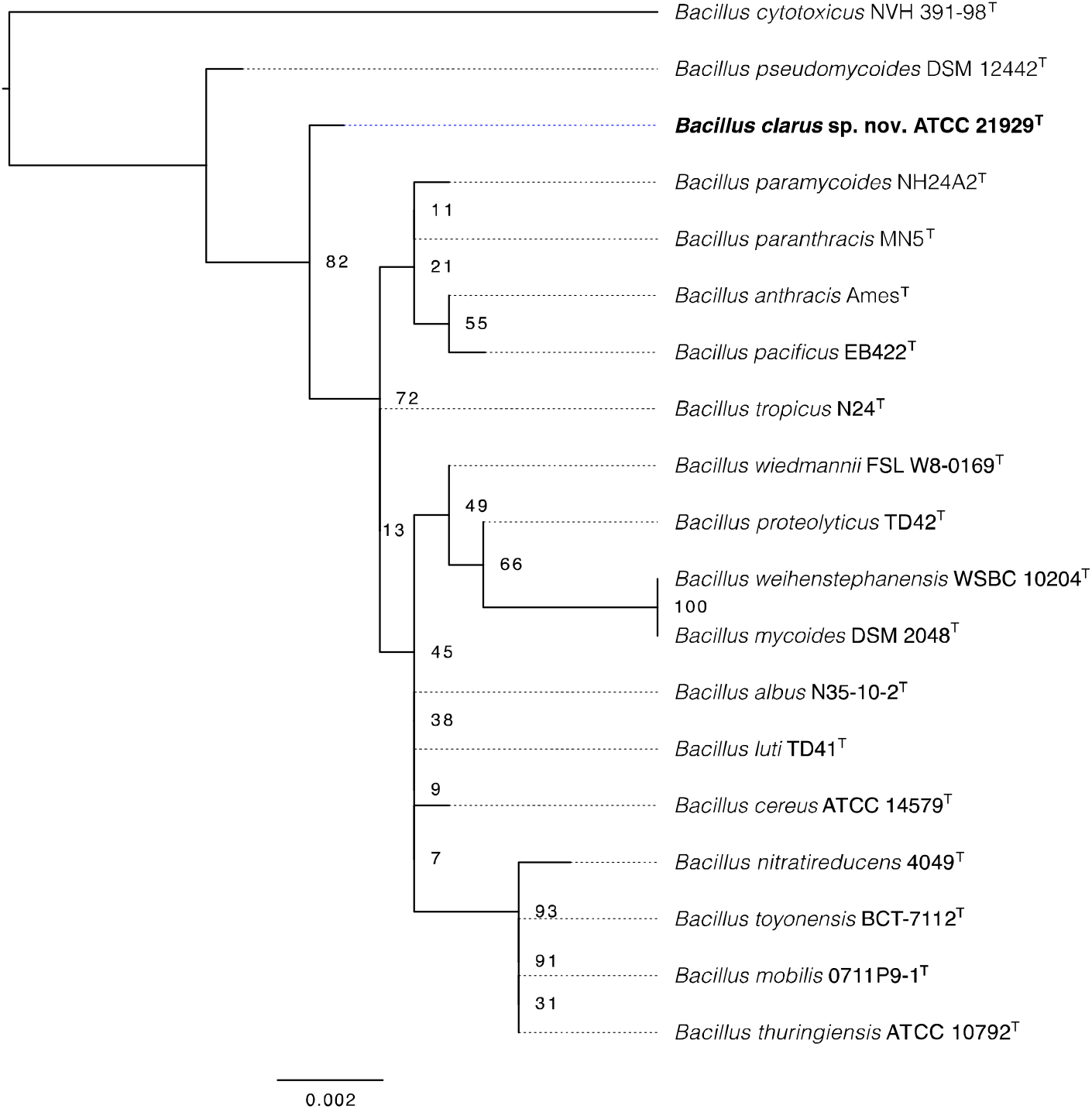
Maximum likelihood phylogeny constructed using 16S rRNA gene sequences of the 18 currently-recognized *B. cereus* group species and novel *B. cereus* group species *B. clarus* ATCC 21929^T^ (boldfaced tip label). The tree is rooted at the midpoint, and node labels correspond to bootstrap support percentages using 1,000 replicates.

**Table 1.** List of *B. cereus* group genomes used in this study.

| Species | Strain | Species Status<sup>a</sup> | ResSeq Accession | *panC* Clade<sup>b</sup> | MLST ST<sup>c</sup> | *rpoB* AT<sup>d</sup> | FastANI ANI<sup>e</sup> | JSpeciesWS<br>ANIb (%<br>aligned)<sup>f</sup> |
| --- | --- | --- | --- | --- | --- | --- | --- | --- |
| *B. clarus* sp. nov. | ATCC 21929 | Novel | GCF_000746925.1 | NA | NA | NA | 100 | 100 |
| *B. albus* | N35-10-2 | Published | GCF_001884185.1 | II | 775 | NA | 85.8323 | 84.84 (64.68) |
| *B. anthracis* | Ames | Published | GCF_000007845.1 | III | 1 | NA | 86.0863 | 85.01 (61.66) |
|  | ATCC 14578 |  |  |  |  |  |  |  |
| *B. cereus* s.s. | ATCC 14579 | Published | GCF_000007825.1 | IV | 921 | 158 | 86.1037 | 84.84 (63.66) |
| *B. cytotoxicus* | NVH 391-98 | Published | GCF_000017425.1 | VII | NA | NA | 83.1993 | 81.36 (48.48) |
| *B. luti* | TD41 | Published | GCF_001884105.1 | II | 764 | NA | 86.1352 | 85.09 (60.76) |
| *B. mobilis* | 0711P9-1 | Published | GCF_001884045.1 | II | 784 | 194 | 86.0059 | 84.81 (61.23) |
| *B. mycoides* | DSM 2048 | Published | GCF_000003925.1 | VI | 116 | 3 | 86.9787 | 86.31 (64.88) |
| *B. nitratireducens* | 4049 | Published | GCF_001884135.1 | VI | 769 | NA | 86.8859 | 86.17 (64.12) |
| *B. pacificus* | EB422 | Published | GCF_001884025.1 | II | 32 | 380 | 85.9859 | 84.92 (61.75) |
| *B. paramycoides* | NH24A2 | Published | GCF_001884235.1 | VI | 780 | NA | 87.3328 | 86.70 (64.31) |
| *B. paranthracis* | Mn5 | Published | GCF_001883995.1 | II | 761 | 125 | 85.9614 | 84.90 (62.38) |
| *B. proteolyticus* | TD42 | Published | GCF_001884065.1 | VI | 765 | NA | 86.7637 | 86.23 (65.23) |
| *B. pseudomycoides* | DSM 12442 | Published | GCF_000161455.1 | I | 83 | 148 | 84.8949 | 83.90 (59.94) |
| *B. thuringiensis* | ATCC 10792 | Published | GCF_002119445.1 | IV | 10 | NA | 85.9932 | 84.83 (64.90) |
| *B. toyonensis* | BCT-7112 | Published | GCF_000496285.1 | V | 111 | 129 | 86.563 | 85.38 (61.22) |
| *B. tropicus* | N24 | Published | GCF_001884035.1 | II | 771 | NA | 85.9761 | 85.02 (62.14) |
| *B. weihenstephanensis* | WSBC 10204 | Published | GCF_000775975.1 | VI | 196 | 3 | 86.9523 | 86.14 (64.40) |
|  | DMS 11821 |  |  |  |  |  |  |  |
| *B. wiedmannii* | FSL W8-0169 | Published | GCF_001583695.1 | II | 1081 | 61 | 86.1474 | 85.00 (63.07) |
| *B. bingmayongensis* | FJAT-1383 | Effective | GCF_000299035.1 | NA | 763 | NA | 87.7067 | 83.67 (58.92) |
| *B. gaemokensis* | JCM 15801 | Effective | GCF_000712615.1 | NA | 768 | NA | 84.8374 | 84.08 (60.40) |
|  | BL3-6 |  |  |  |  |  |  |  |
| *B. manliponensis* | JCM 15802 | Effective | GCF_000712595.1 | NA | NA | NA | 80.4405 | 77.01 (43.10) |
|  | BL4-6 |  |  |  |  |  |  |  |

<sup>a</sup>Novel (proposed in this paper), Published (one of the 18 currently-recognized published and validated *B. cereus* group species), or Effective (previously proposed in a publication as a potential member of the *B. cereus* group, but not validated/published as an official species).
<sup>b</sup>*panC* clade assigned *in silico* using BTyper version 2.2.2 using traditional 7-clade scheme proposed by Guinebretiere, et al. [17].
<sup>c</sup>Multi-locus sequence typing sequence type; assigned *in silico* with BTyper version 2.2.2 [18].
<sup>d</sup>*rpoB* allelic type; assigned *in silico* with BTyper version 2.2.2 [18].
<sup>e</sup>Average nucleotide identity (ANI) calculated using FastANI [19]; proposed novel species *B. mycoides* str. BHP was used as a query and the listed genome was used as a reference.
<sup>f</sup>Average nucleotide identity blast (ANIb) calculated using the JSpecies Web Server (JSpeciesWS [26]), with percent of the genome aligned in parentheses; proposed novel species *B. mycoides* str. BHP was used as a query and the listed genome was used as a reference.

BTyper version 2.2.2 was additionally used to perform *in silico* virulence gene detection, *panC* clade assignment, 7-gene multi-locus sequence typing (MLST), and *rpoB* allelic typing using the assembled genome of *B. clarus* ATCC 21929^T^. FastANI [19] was initially used to compute ANI values between the genome of *B. clarus* ATCC 21929^T^ and the genomes representative of the 18 currently-recognized *B. cereus* group species (Table 1), as well as those of the three effective *B. cereus* group species (Table 1). All ANI values produced by FastANI were < 95, indicating that *B. clarus* ATCC 21929^T^ meets the genomic criteria for a new species. To confirm this using a blast-based ANI metric (ANIb), pyani version 0.2.7 [25] was used to calculate pairwise ANIb values between proposed novel species *B. clarus* ATCC 21929^T^, representative genomes of 18 published *B. cereus* group species, and the type strains of 3 unpublished effective *B. cereus* group species. Furthermore, JSpeciesWS [26] was used to confirm that the pairwise ANIb results produced by pyani were below the 95 ANIb threshold for bacterial species (Table 1).

Core single nucleotide polymorphisms (SNPs) were identified in 22 *B. cereus* group genomes, including the proposed novel species *B. clarus* ATCC 21929^T^, representative genomes of 18 published *B. cereus* group species, and the type strains of three unpublished effective *B. cereus* group species (Table 1), using kSNP version 3.1 [27, 28] with the optimal *k*-mer size determined by Kchooser (*k* = 19). A maximum likelihood (ML) phylogeny was constructed using the resulting core SNPs and RAxML version 8.2.4 [24], using the GTR nucleotide substitution model under the Gamma model of rate heterogeneity, a Lewis ascertainment bias correction [29], and 1,000 bootstrap replicates. FigTree version 1.4.3 (http://tree.bio.ed.ac.uk/software/figtree/) and Graphics for Mac 3.1. were used to annotate the phylogeny (Fig. 2).

**Figure 2.**
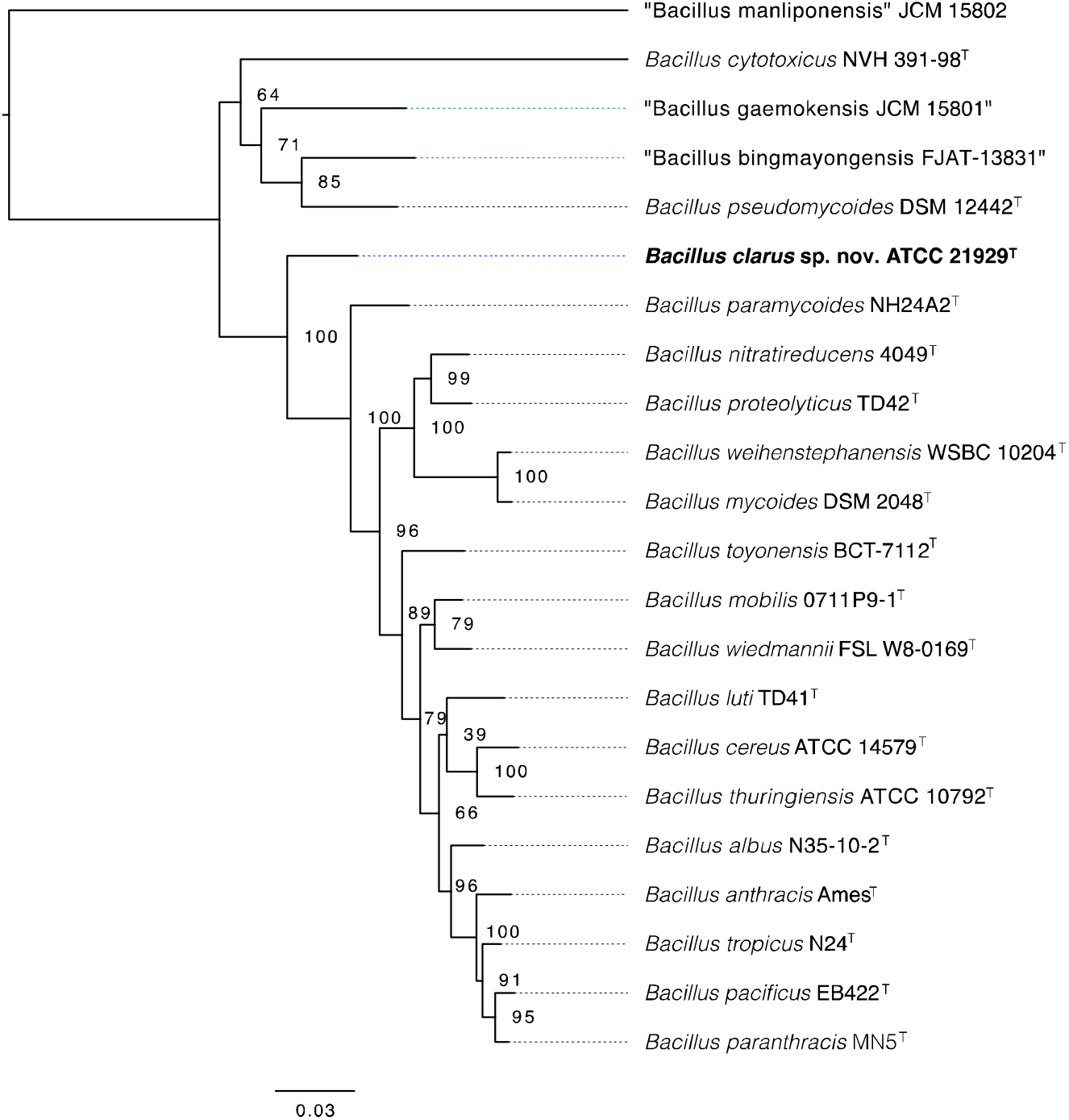
Maximum likelihood phylogeny constructed using core SNPs identified in 18 currently-recognized *B. cereus* group species, 3 proposed effective *B. cereus* group species (tip labels in quotation marks), and novel *B. cereus* group species *B. clarus* ATCC 21929^T^ (boldfaced tip label). The tree is rooted at the midpoint, and node labels correspond to bootstrap support percentages using 1,000 replicates.

Novel *B. cereus* group species *B. clarus* ATCC 21929^T^ could not be assigned to any one of the 7 *panC* phylogenetic clades described by Guinebretiere, et al. [17] based on its *panC* sequence (Table 1). Additionally, it could not be assigned to any known sequence type or allelic type using *in silico* MLST or *rpoB* allelic typing, respectively (Table 1). The strain had MLST allelic types *glp* 97, *ilv*82, *pta* 83, *pur* 82, *pyc* 72, *tpi* 76, and a new *gmk* allele *(gmk* 179) was defined by submission to the PubMLST database, resulting in a new ST 1834. The isolate has been deposited in PubMLST Isolates database under id 2468. The closest match to the *B. clarus* ATCC 21929^T^ *rpoB* sequence in the *rpoB* database of Food Microbe Tracker was allelic type 342 (AT0342): the two alleles shared 95.1% identity and 99.5% query coverage [30]. The 16S rDNA sequence of *B. clarus* ATCC 21929^T^ most closely resembled that of the *B. tropicus* type strain (Figure 1), which it matched with 99.8% and 100% nucleotide and coverage, respectively. However, based on both core SNPs detected in all valid and effective *B. cereus* group species, as well as its ANIb metric, the novel species most closely resembled *B. paramycoides* (Figs. 2 and S1). The pairwise ANIb values for *B. clarus* ATCC 21929^T^ and *B. paramycoides*NH24A2^T^ were 87.33 % and 86.70%, as determined by FastANI and JSpeciesWS (Table 1). The DDH value for these same two strains was 34.1%, with 0.48% probability for DDH being >70% (Table S1), providing strong evidence for a new genomospecies. Genes coding for diarrheal enterotoxin Hbl (*hblABCD*) were detected in the genome of *B. clarus* ATCC 21929^T^.

## Phenotypic characteristics

All phenotypic tests were conducted using *B. cereus* ATCC 14579^T^ as a control strain. *B. clarus* ATCC 21929^T^ was confirmed as weakly-positive for production of Hbl and negative for production of Nhe using the Duopath Cereus Enterotoxins kit (Merck) when the culture was grown to the stationary phase at 32°C without shaking. At 37°C, *B. clarus* ATCC 21929^T^ did not produce Hbl nor Nhe when grown in the same conditions. *B. cereus* s.s. strain ATCC 14579^T^, which was used as a control, was positive for production of both toxins when grown at both temperatures without shaking. The cytotoxic potential of *B. clarus* ATCC 21929^T^ was assessed in a 96-well microtiter plate by incubating 12 replicates of confluent HeLa cells with 5% v/v bacterial supernatant (bacteria grown at 37°C) for 15 min, followed by addition of 10 μl of WST-1 dye solution (Roche) and further 25-min incubation [2]. The final absorbance was determined by subtracting the absorbance values measured at 690 nm from those measured at 450 nm. Percent viability was determined relative to cells treated with 5% v/v Brain Heart Infusion (BHI; negative cytotoxicity control). 0.05% Triton X-100 was used as a positive cytotoxicity control (Figure 3) [2].

**Figure 3.**
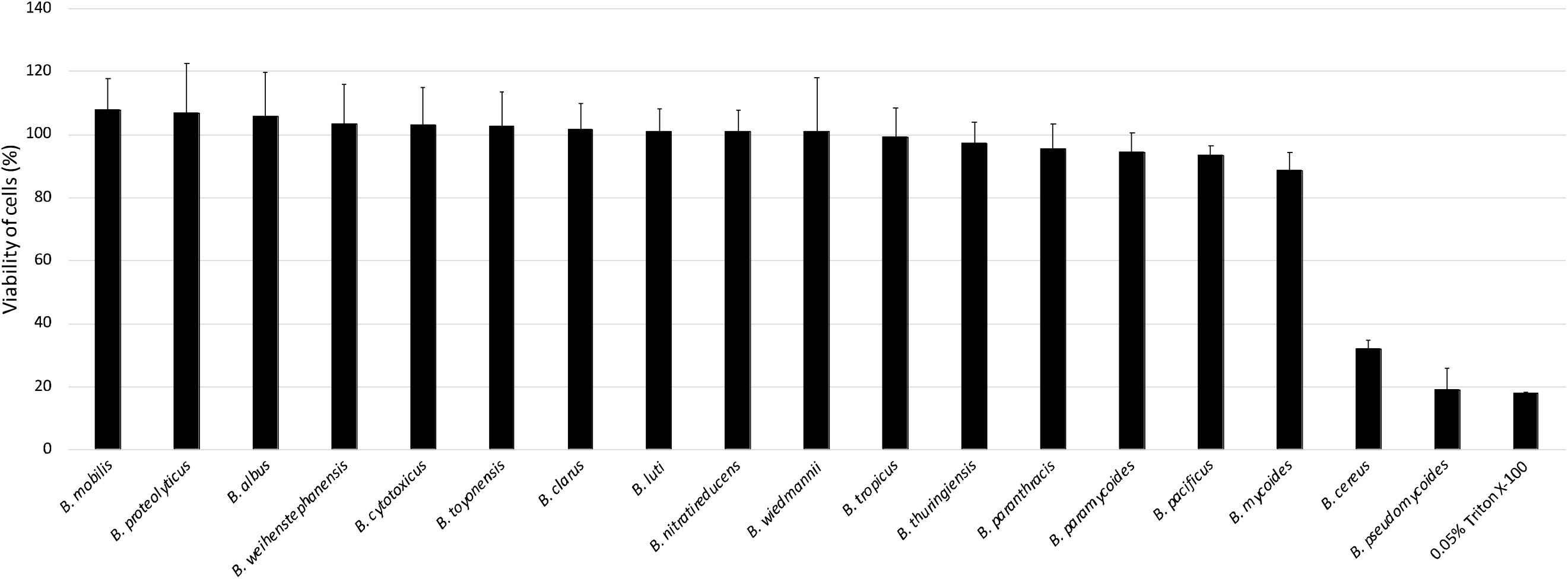
Cytotoxicity of *B. clarus* ATCC 21929^T^ and 17 other valid *B. cereus* group species type strains. Percentage viability of HeLa cells when treated with supernatants of *B. clarus* ATCC 21929^T^ and 17 other valid *B. cereus* group type strains as determined by the WST-1 assay [2]. The columns represent the mean viabilities and the error bars represent standard deviations for 12 technical replicates.

Gram staining was utilized for the preliminary identification of *B. clarus* ATCC 21929^T^. Cells stained Gram-positive and were approximately 3 μm long. Additionally, the morphology of the bacteria grown overnight in BHI broth at 32°C and stained with 2% uranyl acetate negative stain was observed by transmission electron microscopy (Figure S1). *B. clarus* ATCC 21929^T^ was hemolytic, as confirmed by zones of clearance after streaking a loopful of 24-hour culture suspension onto blood agar and incubating it at 35°C for 24 h. *B. clarus* ATCC 21929^T^ was oxidase-negative, as confirmed using the Oxidrop reagent (Hardy Diagnostics). It was able to hydrolyze starch and casein at 32°C after 72 hours of incubation, indicating that the strain possesses both amylase and caseinase activity. Assays were conducted following protocols described in Bergey’s manual [16]. The ability of *B. clarus* ATCC 21929^T^ to grow in anaerobic conditions was tested by inoculating anaerobic agar with an overnight culture and incubating it in a jar with anaerobic gas pack at 30°C for 7 days; visible growth was observed after 3 days of incubation. The motility was examined by preparing a Motility Test Medium according to the Bacteriological Analytical Manual (BAM) [31], stab-inoculating overnight culture, and incubating it for 24 h at 32°C. The strain’s ability to grow at different temperatures (4, 7, 10, 15, 20, 25, 30, 37, 40, 43, 45, and 55°C) was studied by streaking individual well isolated colonies on BHI agar plates, in triplicates, and incubating them up to 3 weeks or until growth was observed (Table 1) [16]. The ability of *B. clarus* ATCC 21929^T^ to grow at different pH was confirmed by inoculating 10 μl of an overnight culture into BHI broths adjusted to pH 3–11 using appropriate buffers, in triplicate. Citrate buffer was used to supplement BHI adjusted to pH 3, 4, and 5, phosphate buffer was added to BHI adjusted to pH 6,7, and 8, and CAPS buffer was added in BHI adjusted to pH 9,10, and 11. Inoculated BHI tubes were incubated at 30°C for 14 days or until growth was observed based on turbidity. The ability to grow at different concentrations of NaCl was determined by supplementing TSB broth with 0, 0.5, 1, 2, 3, 5, 7, 9, 12, and 15% of NaCl. Tubes were inoculated with 10 μl of an overnight culture, in triplicate, and incubated at 32°C for 14 days or until growth was confirmed based on turbidity. Results of the phenotypic tests described above are reported in Table 2. The fatty acid composition was determined by MID Inc. for culture grown at their defined standard conditions, on tryptic soy agar at 28°C. API 20E and CH50 biochemical assays (bioMérieux) were performed following the manufacturer’s instructions at 32°C (Table 3).

**Table 2.**
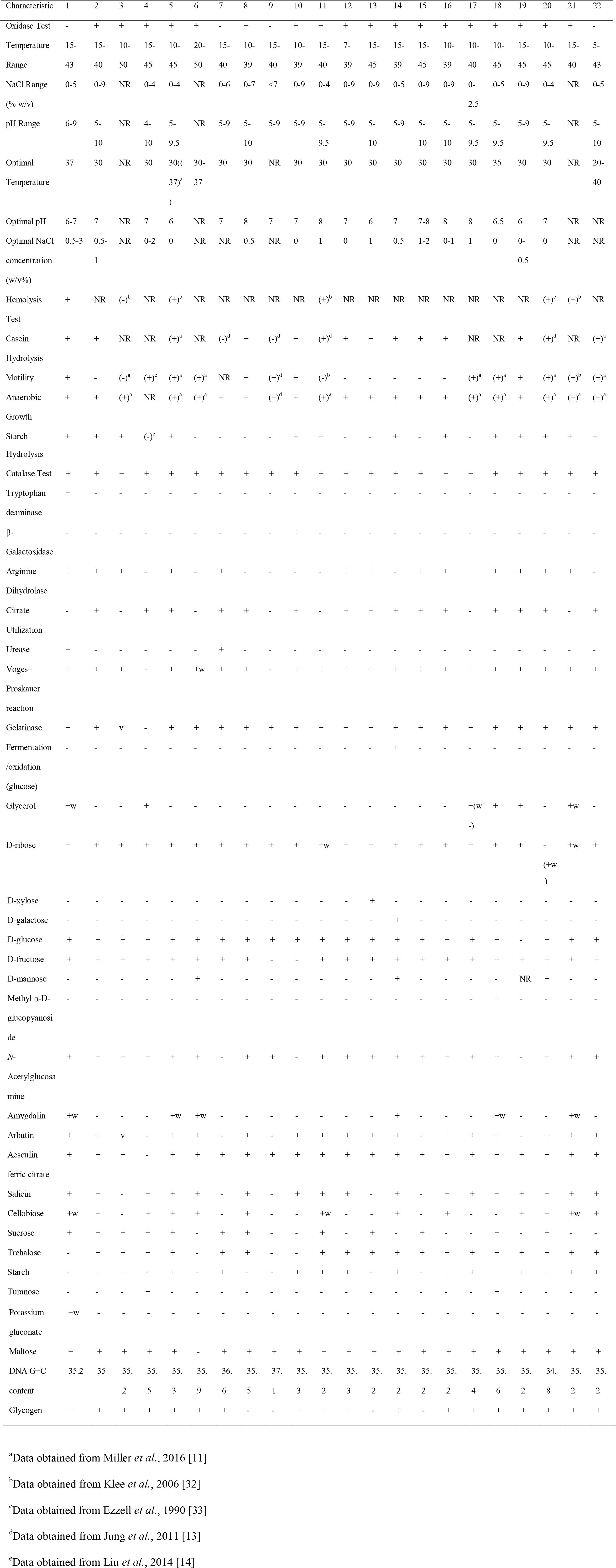
Phenotypic characteristics of *B. clarus* ATCC 21929^T^ and other valid and effective *Bacillus cereus* group species type strains. Species: 1, *B. clarus* ATCC 21929; 2, *B. albus*N35-10-2; 3, *B. anthracis* ATCC 14578; 4, *bingmayogensis* FJAT-13831; 5, *B. cereus* ATCC 14579; 6, *B. cytotoxicus* NVH 391-391; 7, *B. gaemokensis* BL3-3; 8, *B. luti* TD41; 9, *B. manliponensis* BL4-4; 10, *B. mobilis* 0711P9-9; 11, *B. mycoides* DSM 2048; 12, *B. nitratireducens* 4049; 13, *B. pacificus*EB422; 14, *B. paramycoides* NH24A2; 15, *B. paranthracis* Mn5; 16, *B. proteolyticus* TD42; 17, *B. pseudomycoides* DSM 12442; 18, *B. toyonensis*BCT-7112; 19, *B. tropicus* N24; 20, *B. thuringiensis* ATCC 10792; 21, *B. weihenstephanesis* DSM 11821; 22, *B. wiedmannii* FSL W-0169. The data for strain ATCC 21929 were produced in this study. All other data were obtained from Liu *et al*, 2017 [4] unless specified otherwise in footnotes. In the API 20E tests, all strains were negative for lysine decarboxylase, ornithine decarboxylase, H_2_S production, indole production, mannitol, inositol, sorbitol, rhamnose, melibiose, and arabinose. In the API 50CHB tests, all strains were negative for erythritol, D-arabinose, L-arabinose, L-xylose, D-adonitol, methyl *β*-D-xylopyranoside, L-sorbose, L-rhamnose, dulcitol, inositol, D-mannitol, D-sorbitol, methyl *α*-D-mannopyranoside, lactose, melibose, inulin, melezitose, raffinose, xylitol, gentiobiose, D-lyxose, D-tagatose, D-fucose, L-fucose, D-arabitol, L-arabitol, potassium 2-ketogluconate and potassium 5-ketogluconate. -, negative; -w, weakly negative; +, positive; +w, weakly positive; NR, not reported.

**Table 3:**
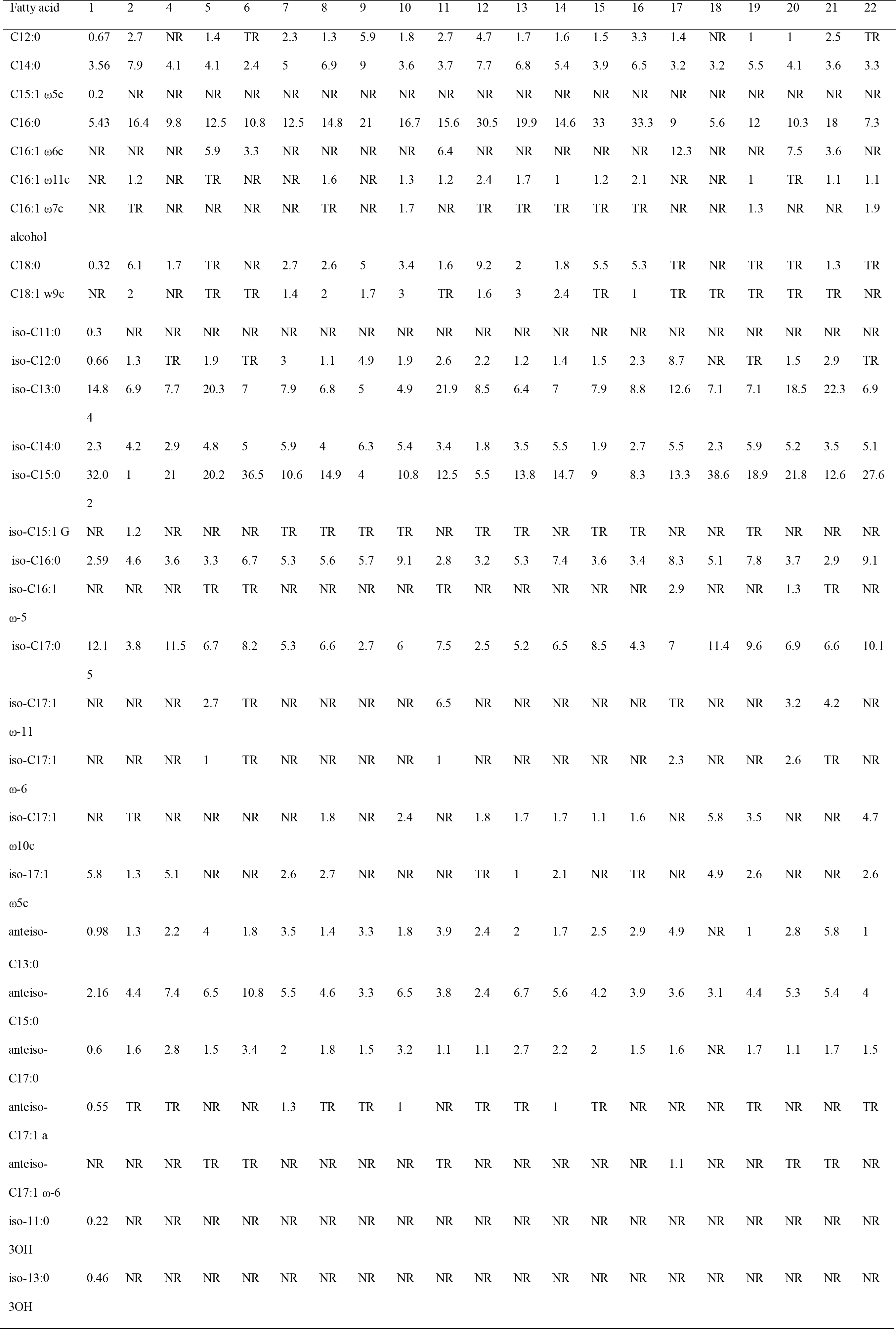
Fatty acid composition of *B. clarus* ATCC 21929^T^ and other valid and effective *Bacillus cereus* group species type strains. Species: 1, *B. clarus* ATCC 21929; 2, *B. albus*N35-10-2; 3, *B. anthracis* ATCC 14578 (no data to show); 4, *bingmayogensis* FJAT-13831; 5, *B. cereus* ATCC 14579; 6, *B. cytotoxicus* NVH 391-391; 7, *B. gaemokensis*BL3-3; 8, *B. luti* TD41; 9, *B. manliponensis* BL4-6; 10, *B. mobilis* 0711P9-9; 11, *B. mycoides* DSM 2048; 12, *B. nitratireducens* 4049; 13, *B. pacificus*EB422; 14, *B. paramycoides* NH24A2; 15, *B. paranthracis* Mn5; 16, *B. proteolyticus* TD42; 17, *B. pseudomycoides*DSM 12442; 18, *B. toyonensis*BCT-7112; 19, *B. tropicus*N24; 20, *B. thuringiensis* ATCC 10792; 21, *B. weihenstephanensis* DSM 11821; 22, *B. wiedmannii* FSL W8-0169. The data for strain ATCC 21929 was produced in this study. All other data were obtained from Liu *et al*, 2017 [4]. NR: not reported; TR: trace amount.

## Description of *Bacillus clarus* sp.nov

*Bacillus clarus* (cla’rus. L. masc. adj. *clarus* clear).

Cells stained as Gram-positive and displayed a long rod-like appearance, 3 μm in length. *B. clarus* ATCC 21929^T^ is highly motile, oxidase negative, hemolytic, possesses amylase and caseinase activity, can reach stationary phase in 16 hours at a growth temperature of 32°C in BHI, and can grow in aerobic and anaerobic conditions. *B. clarus* ATCC 21929^T^ can grow at pH 6-9, temperatures ranging from 15°C-43°C, and salt concentrations of 0-5%. The optimum conditions of growth are 6-9, 37°C, and 0.5-3% respectively. *B. clarus* ATCC 21929^T^ is weakly positive hemolysin BL (HBL) production at a temperature of 32°C, as indicated by faint bands in Duopath Enterotoxins test, but does not reduce the metabolic activity of HeLa cells at tested conditions. The fatty acid that was highest in abundance was iso-C15:0. Amongst the least abundant fatty acids were C15:1 ω5c and iso-11:0 3OH. The latter two fatty acids, along with iso-13:0 3OH were not found to be reported for any of the other *B. cereus* group type strains (Table 3). Unique characteristics of *B. clarus* ATCC 21929^T^ include a higher abundance of iso-C17:0, lower abundance of iso-C16:0 fatty acids, and the ability to grow optimally even at 3% NaCl concentration. *B. clarus* ATCC 21929^T^ is also oxidase negative, which is a trait shared by only *B. wiedmannii, B. gaemokensis*, and *B. manliponensis* of the *B. cereus* group (Table 2).

## Authors Statements

There are no conflicts of interest and all funding provided to support this study is listed in Acknowledgments. No Ethical Committee approvals were needed for this study.

## Acknowledgements

The authors would like to thank to Ryan Michael Gaboy and bioMérieux for a generous support of this project through donation of API 20E and CH50 assay kits. M. A. M was supported by the USDA-funded REEU project “Bugs in my Food: Research and Professional Development in Food Safety for Undergraduates from Non-Land Grant Institutions” (USDA-NIFA 2017-67032-26022), and L. M. C by the National Science Foundation Graduate Research Fellowship Program under grant no. DGE-1144153. J. K. and E. G. D. were supported by the USDA National Institute of Food and Agriculture Hatch Appropriations under projects #PEN04646 and #PEN04522 and accessions #1015787 and #0233376, respectively. L. X. and sequencing were supported by the U.S. Food and Drug Administration grant number 1U18FD006222-01 in support of the GenomeTrakr in Pennsylvania.

## Abbreviations

MLST: Multi-Locus Sequence Type
ANIb: Average Nucleotide Identity BLAST
BAM: Bacteriological Analytical Manual
DDH: DNA-DNA hybridization
GTR: generalized time-reversible model
ML: maximum likelihood
MLST: multilocus sequence typing
SNP: single nucleotide polymorphism
SRA: Sequence Read Archive
WGS: whole genome sequence

