## Supplementary Materials for "*Bacillus clarus* sp. nov. is a new *Bacillus cereus* group species isolated from soil"

**Journal:** International Journal of Systematic and Evolutionary Microbiology

**Fig. S1. Heatmap of pairwise average nucleotide identity values computed using the BLAST algorithm (ANIb).** Rows and columns correspond to 18 currently-recognized *B. cereus* group species, 3 effective *B. cereus* group species, and the novel *B. cereus* group species type strain *B. clarus* str. ATCC 21929^T^ (=PS00077A^T^ = PS00077B^T^ = PSU-0922^T^ = BHP^T^).
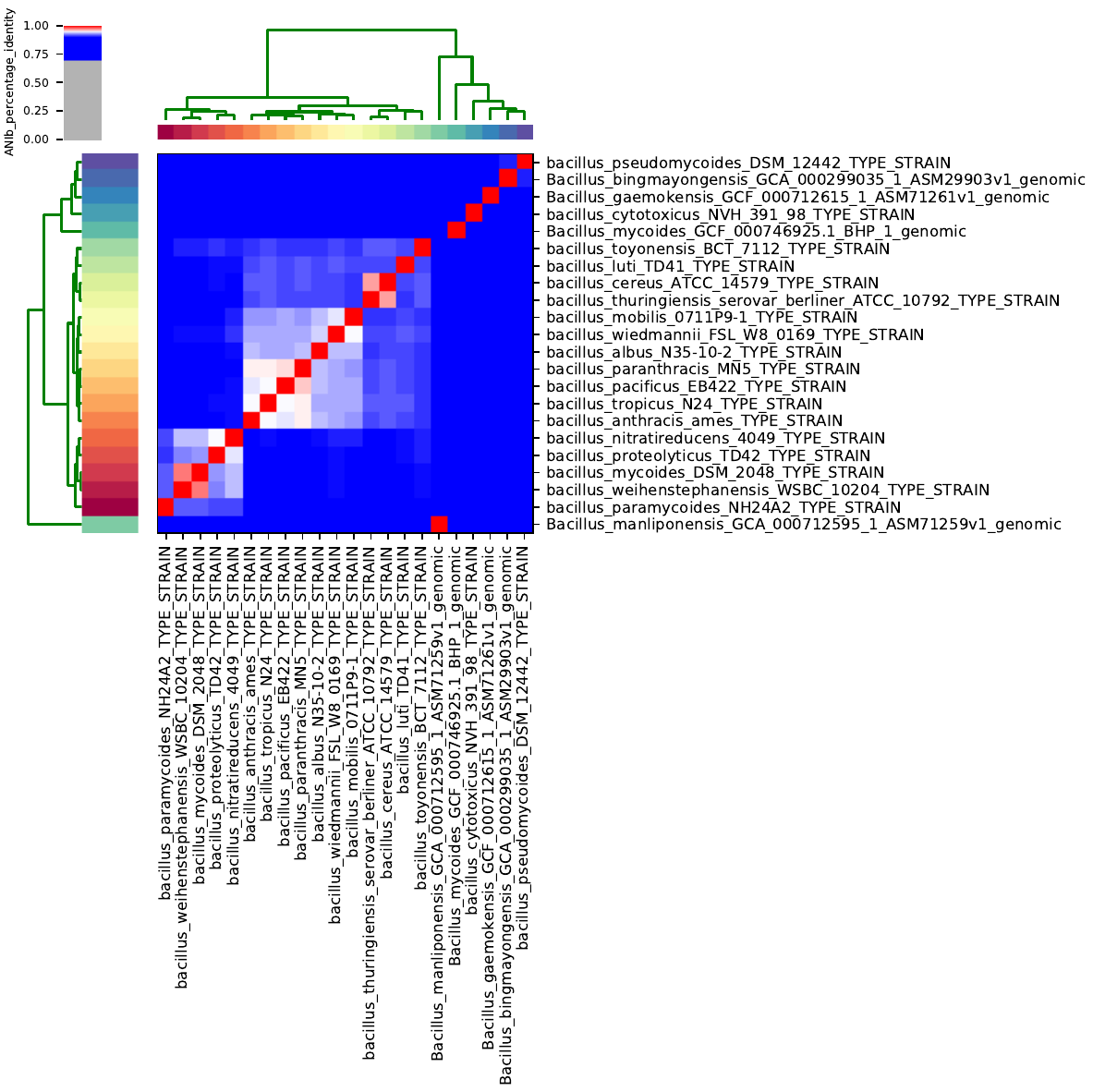

**Fig. S2. Transmission electron microscopy image of *B. clarus* ATCC 21929^T^.** Image was obtained using 2% uranyl acetate staining.

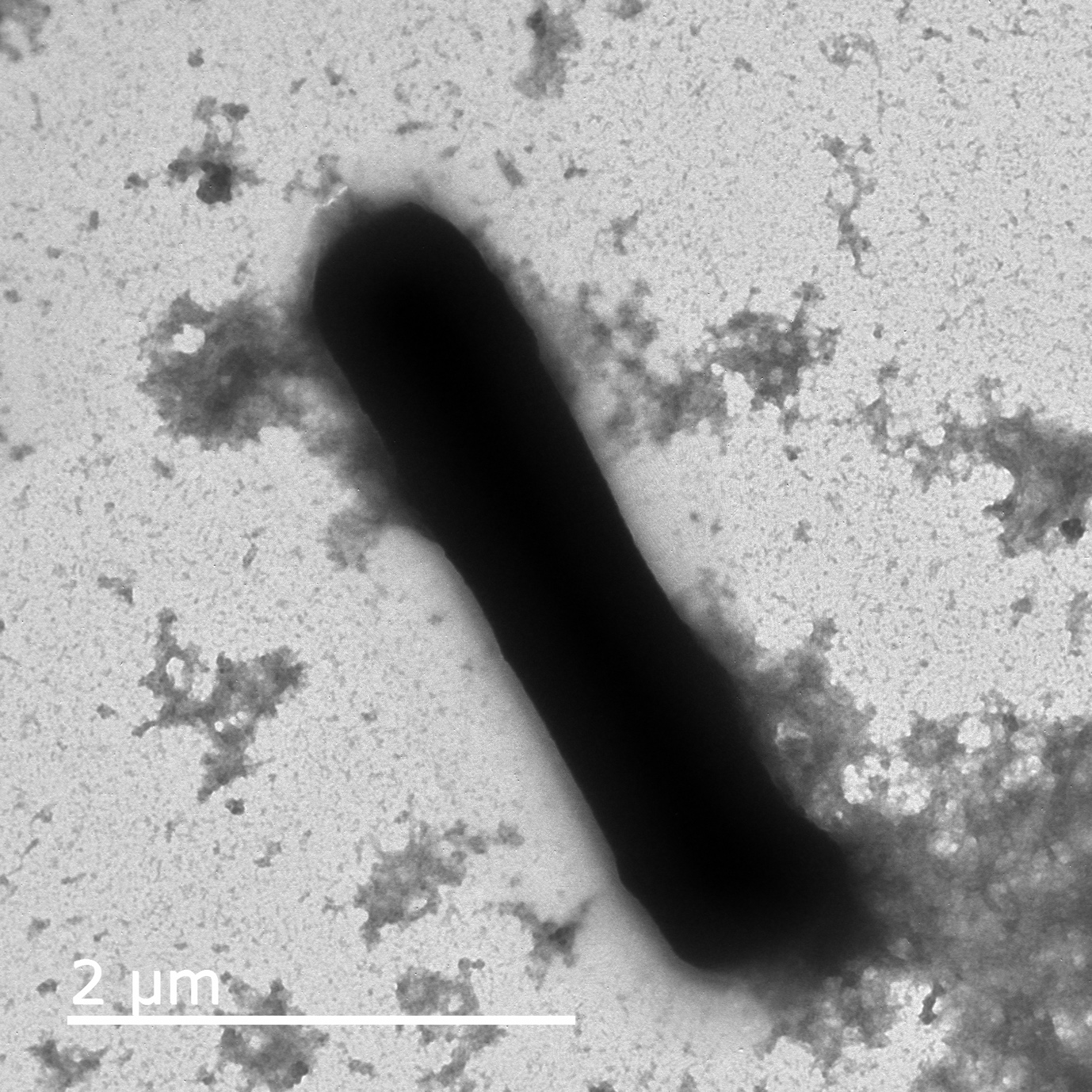

**Table S1.** Predicted DNA-DNA hybridization (DDH) between *B. clarus* ATCC 21929^T^ and representative strains of other *B. cereus* group species.

| Query genome | Reference genome | DDH^a^ | Model C.I.b | Distance | Prob. DDH >= 70% | G+C difference |
| --- | --- | --- | --- | --- | --- | --- |
| *B. clarus* sp. nov str. ATCC 21929 | *B. albus* N35-10-2 | 30.7 | [28.4 - 33.3%] | 0.1381 | 0.14 | 0.37 |
| *B. clarus* sp. nov str. ATCC 21929 | *B. anthracis* Ames | 31 | [28.6 - 33.5%] | 0.137 | 0.16 | 0.05 |
| *B. clarus* sp. nov str. ATCC 21929 | *B. bingmayongensis* FJAT-13831 | 29.5 | [27.1 - 32%] | 0.1449 | 0.08 | 0.17 |
| *B. clarus* sp. nov str. ATCC 21929 | *B. cereus* ATCC 14579 | 30.9 | [28.5 - 33.4%] | 0.1373 | 0.15 | 0.03 |
| *B. clarus* sp. nov str. ATCC 21929 | *B. cytotoxicus* NVH 391-98 | 26.9 | [24.6 - 29.4%] | 0.1607 | 0.03 | 0.54 |
| *B. clarus* sp. nov str. ATCC 21929 | *B. gaemokensis* JCM 15801 | 29.3 | [26.9 - 31.8%] | 0.1459 | 0.08 | 0.4 |
| *B. clarus* sp. nov str. ATCC 21929 | *B. luti* TD41 | 31 | [28.6 - 33.5%] | 0.1368 | 0.16 | 0.13 |
| *B. clarus* sp. nov str. ATCC 21929 | *B. manliponensis* JCM 15802 | 23.3 | [21 - 25.7%] | 0.188 | 0 | 0.7 |
| *B. clarus* sp. nov str. ATCC 21929 | *B. mobilis* 0711P9-1 | 30.7 | [28.3 - 33.2%] | 0.1383 | 0.14 | 0.02 |
| *B. clarus* sp. nov str. ATCC 21929 | *B. mycoides* DSM 2048 | 33.2 | [30.8 - 35.7%] | 0.126 | 0.36 | 0.11 |
| *B. clarus* sp. nov str. ATCC 21929 | *B. nitratireducens* 4049 | 32.6 | [30.2 - 35.1%] | 0.129 | 0.29 | 0.01 |
| *B. clarus* sp. nov str. ATCC 21929 | *B. pacificus* EB422 | 30.6 | [28.2 - 33.1%] | 0.1388 | 0.14 | 0.11 |
| *B. clarus* sp. nov str. ATCC 21929 | *B. paramycoide*s NH24A2 | 34.1 | [31.6 - 36.6%] | 0.1222 | 0.48 | 0.11 |
| *B. clarus* sp. nov str. ATCC 21929 | *B. paranthracis* MN5 | 30.6 | [28.2 - 33.1%] | 0.139 | 0.13 | 0.13 |
| *B. clarus* sp. nov str. ATCC 21929 | *B. proteolyticus* TD42 | 32.7 | [30.3 - 35.2%] | 0.1284 | 0.3 | 0.17 |
| *B. clarus* sp. nov str. ATCC 21929 | *B. pseudomycoides* DSM 12442 | 30.2 | [27.8 - 32.7%] | 0.1411 | 0.11 | 0.05 |
| *B. clarus* sp. nov str. ATCC 21929 | *B. thuringiensis* serovar berliner ATCC 10792 | 30.7 | [28.3 - 33.2%] | 0.1386 | 0.14 | 0.5 |
| *B. clarus* sp. nov str. ATCC 21929 | *B. toyonensis* BCT-7112 | 32.1 | [29.7 - 34.6%] | 0.1313 | 0.24 | 0.23 |
| *B. clarus* sp. nov str. ATCC 21929 | *B. tropicus* N24 | 30.7 | [28.3 - 33.2%] | 0.1383 | 0.14 | 0.1 |
| *B. clarus* sp. nov str. ATCC 21929 | *B. weihenstephanensis* WSBC 10204 | 32.7 | [30.2 - 35.2%] | 0.1286 | 0.3 | 0.08 |
| *B. clarus* sp. nov str. ATCC 21929 | *B. wiedmannii* FSL W8-0169 | 31 | [28.6 - 33.5%] | 0.1368 | 0.16 | 0.13 |

**^a^** DDH, predicted DNA-DNA hybridization values calculated using formula 2 of the Genome-to-Genome Distance Calculator (GGDC 2.1; Meier-Kolthoff et al., 2013), which is recommended for draft genome analysis.

^b^ C.I., estimated confidence interval.
